# Morphological bias of the MNI152 brain

**DOI:** 10.64898/2026.01.24.701532

**Authors:** Yishai Valter, Yu Huang, Niranjan Khadka, Marom Bikson, Abhishek Datta

## Abstract

The MNI152 template is widely treated as a representative average brain in neuroimaging, computational modeling, and neuromodulation research, yet its fidelity to true population morphology has not been systematically evaluated. In this study, we compared the MNI152 template to 430 individual MRI scans from a publicly available dataset spanning Asian, Black, and White participants. We additionally generated an alternative template using ANTsPy to assess whether a modern diffeomorphic approach yields a more anatomically representative average.

We conducted affine registrations and deformation-based morphometry to detect and quantify gross morphological differences as well as local voxel-level deformations. Across all racial groups, the MNI152 template required consistent global contraction, and its Jacobian fields revealed spatially heterogeneous deformations, indicating systematic mismatches in both size and shape. In contrast, the ANTsPy template showed mean scaling and deformation values near 1.0, reflecting closer correspondence to real human anatomy.

These findings demonstrate that the MNI152 template does not accurately represent the morphology of the studied population and that linear registration alone cannot correct its inherent biases. Reliance on MNI152 may therefore introduce unintended distortions in applications requiring anatomically realistic head models. More robust, unbiased template-generation pipelines, and potentially demographic-specific templates, may be necessary for improved accuracy.

## Introduction

Standardized brain templates are the foundation of neuroimaging and modeling. They provide a common spatial reference that enables comparisons across individuals, studies, and modalities, and they underpin practically all group-level analyses in MRI, source localization, and computational modeling. Among these templates, the Montreal Neurological Institute (MNI) series has become the de facto coordinate system for human-brain research.

The earliest MNI template (MNI305) was created in the 1990s by manually identifying anatomical landmarks in 305 young adult MRI scans and linearly registering them to approximate Talairach space. Because only linear transformations were used, interindividual differences in cortical folding, cranial proportions, and local curvature accumulated in the group average. This produced a template that was systematically larger than any individual brain, exhibiting a “virtual convolution” (Evans et al., 1993) due to the inherent limitations of early registration methods.

The subsequent MNI152 (ICBM152) template improved the image quality by averaging 152 higher-resolution scans; however, it inherited the global morphology of the original MNI305 space (Mazziotta et al., 1995, 2001a, 2001b). The Colin27 template (Holmes et al., 1998) and nonlinear MNI152 templates (Evans et al., 2012) provided sharper and more accurate anatomical detail but retained the global morphology established by the earlier linear templates. Extended-coverage models, such as the New York Head (Huang et al., 2016) and the Big Field of View template (Kreilkamp et al., 2023), also preserved the MNI152 size and shape.

Prior work has shown that MNI152 is larger than the Talairach atlas and several race-specific templates, the including Chinese, Indian, and Korean population averages (Lancaster et al., 2007; Tang et al., 2010; Lee et al., 2005). Also, studies have shown spatial discrepancies in certain subcortical structures between the MNI152 and individual brains (Giff et al., 2023). However, no study has directly compared MNI152 to a large set of individual subject scans across multiple racial groups, including Caucasians. Such a comparison is essential for determining whether the MNI152 brain truly represents an “average” human brain or whether its morphology primarily reflects the artifacts of template construction.

In this study, we compared the MNI152 brain to a large sample of subject scans to quantify detect and quantify potential anatomical bias. To establish a comparative baseline, we also generated an alternative brain template using ANTsPy (Tustison et al., 2021) and compared it to the same large dataset to determine whether this alternative provided a better approximation of real human brain morphology.

Our findings revealed substantial and spatially nonuniform discrepancies between the MNI152 template and actual subject scans. These discrepancies were notably smaller when comparing the ANTsPy template to the same population, suggesting that reliance on MNI152 may introduce unintended biases in applications requiring anatomically realistic head models. Our work provides support for the use of alternative templates, especially those formed for specific demographic groups.

## Methods

### Dataset and Processing

Structural MR scans were obtained from the Dallas Lifespan Brain Study (Park et al., 2025), a large publicly available dataset hosted on OpenNeuro. The full dataset included 464 unique participant scans spanning a wide adult age range (21-89 years; mean 58.31 ±18.14). For the present analysis, we downloaded T1w volumes from participants who self-identified as Asian (n=14), Black / African-American (n=26), or White / Caucasian (n=395). Participants labeled as mixed race, unknown, or other races were excluded because of their limited sample size. Processing was done using FSL (version 6.0.7.17). Brain extraction was performed on each subject’s scan (FSL’s *bet*) and on the 2mm resolution MNI152 template provided as part of the FSL software package. The segmented brains were visually inspected for segmentation accuracy and six brains were discarded due to failed segmentation (e.g., incomplete extraction, skull remnants).

### Affine transform analysis

Parameter-specific differences can be quantified by registering one image to another using an affine transformation and extracting the scaling factors (*λ*_*x*_ , *λ*_*y*_ , *λ*_*z*_) and shear factors (*h* _*xy*_ , *h*_*xz*_,*h*_*yz*_) from the transformation matrix (Buckner et al., 2004; Ashburner & Friston, 2000). We performed this analysis by registering the MNI152 brain to each subject’s brain-extracted volume (FSL, *flirt*). The transformed brains were visually inspected and eight brains were discarded for failed registration. The resulting affine matrices were decomposed (FSL, *avscale*) to isolate scaling and shear components. A scaling factor >1 indicates expansion and <1 indicates contraction. Absolute values of the shear parameters were converted into degrees using the inverse tangent (θ = *arctan* (|shear|)) with values close to 0 reflecting minimal shear.

### Deformation-based morphometry (DBM)

DBM involves performing a nonlinear registration of one image to another, generating a three-dimensional field of Jacobian determinants (|*J*|). The |*J*| is a scalar value calculated at the voxel level that serves as a localized measure of volume change. A |*J*| > 1 indicates voxel expansion and |*J*| < 1 indicates voxel contraction (Freeborough & Fox, 1998; Gee & Bajcsy, 1999; Ashburner & Friston, 2000; Chung et al., 2001; Leow 2007). By aggregating these voxel-wise maps across an entire cohort, we can quantitatively assess anatomical differences between a template and the study population (Yang et al., 2020). In this study, we deformed the MNI152 brain to each subject’s scan (FSL, *fnirt*). The resulting |*J*| maps were concatenated into a 4D array and averaged across the subject dimension. The final output was a 3D spatial map where values significantly divergent from 1 represent systematic morphological bias relative to the study population.

### Alternative template

To establish a comparative baseline, an alternative brain template was generated using a subset of 50 randomly selected structural scans from the study’s dataset. The ANTsPy library was used to create the alternative template (*ants*.*build_template*) with an unbiased initialization parameter (*initial_template* set to “None”) and configured to optimize over 10 iterations. The brain was segmented from this alternative template (“ANTsPy50”) using the *antspynet*.*brain_extraction* command. As was done with the MNI152 brain, the ANTsPy-generated brain was transformed to each subject brain using a linear affine transformation for calculating scaling and shear parameters, as well as a nonlinear deformation registration for DBM.

## Results

### Linear transform

Across all racial groups, mean scaling factors for the MNI152 brain were consistently below 1.0, indicating that the MNI152 brain must be contracted to match individual anatomy for most subjects (Figure 2). In contrast, registration of the ANTsPy50 brain to each subject’s scan resulted in mean scaling factors close to 1.0. For both templates, means of absolute shear values were <0.05 corresponding to angular distortions of ∼2 degrees or less, demonstrating that shear contributes minimally to interindividual differences in brain morphology.

**Figure 1.**
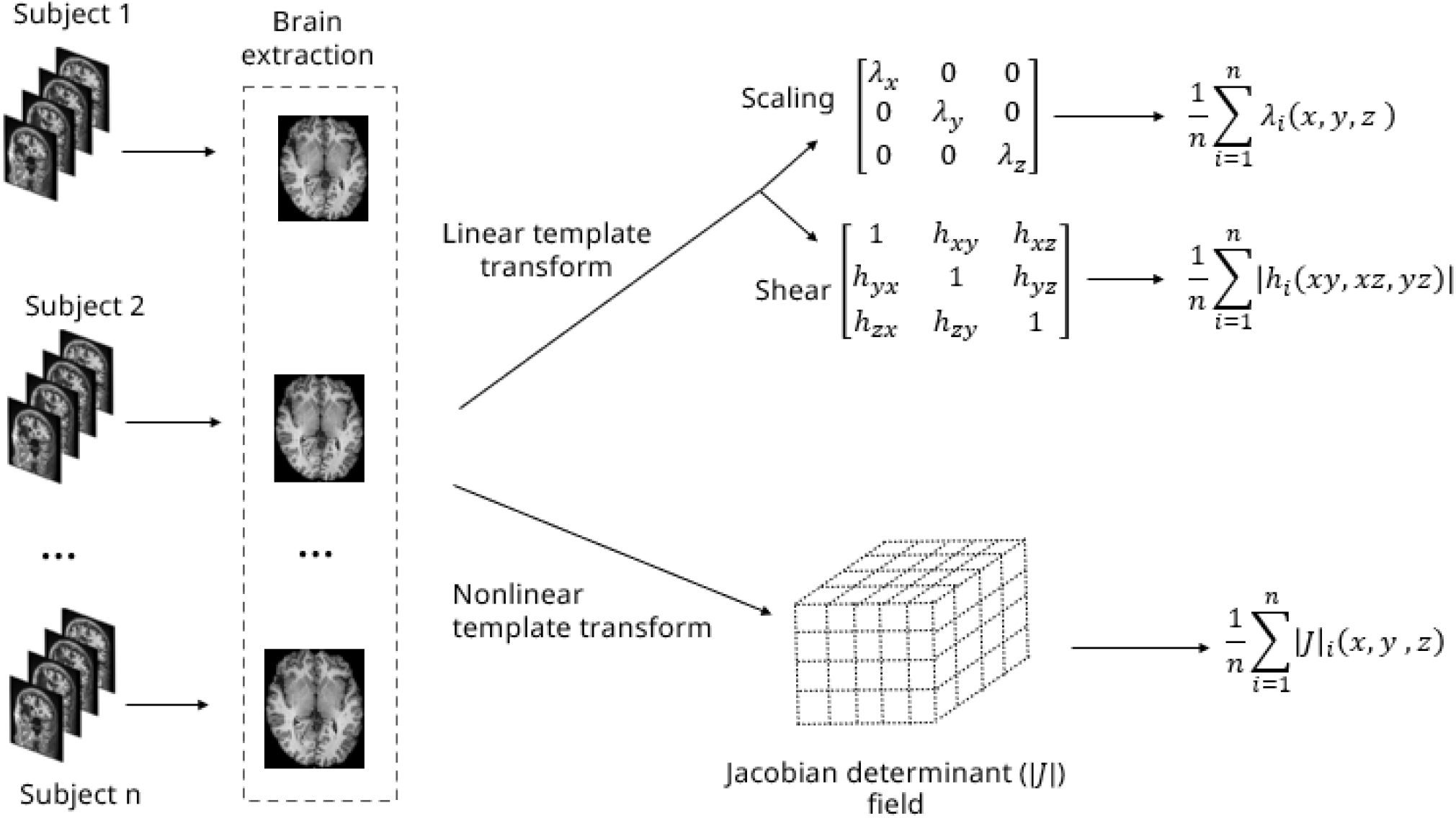
Template-to-subject registration workflow and analysis. Brain tissue was extracted from each subject’s T1-weighted MRI. Two brain templates (MNI152 and ANTsPy-generated brain) were then registered to each subject’s brain using linear (FSL, *flirt*) and nonlinear (FSL, *fnirt*) registration. Affine matrices were decomposed (FSL, *avscale*) to obtain axis-specific scaling factors and shear components. Mean scaling, shear, and voxel-wise Jacobian values were computed to characterize morphological biases in each template.

**Figure 2.**
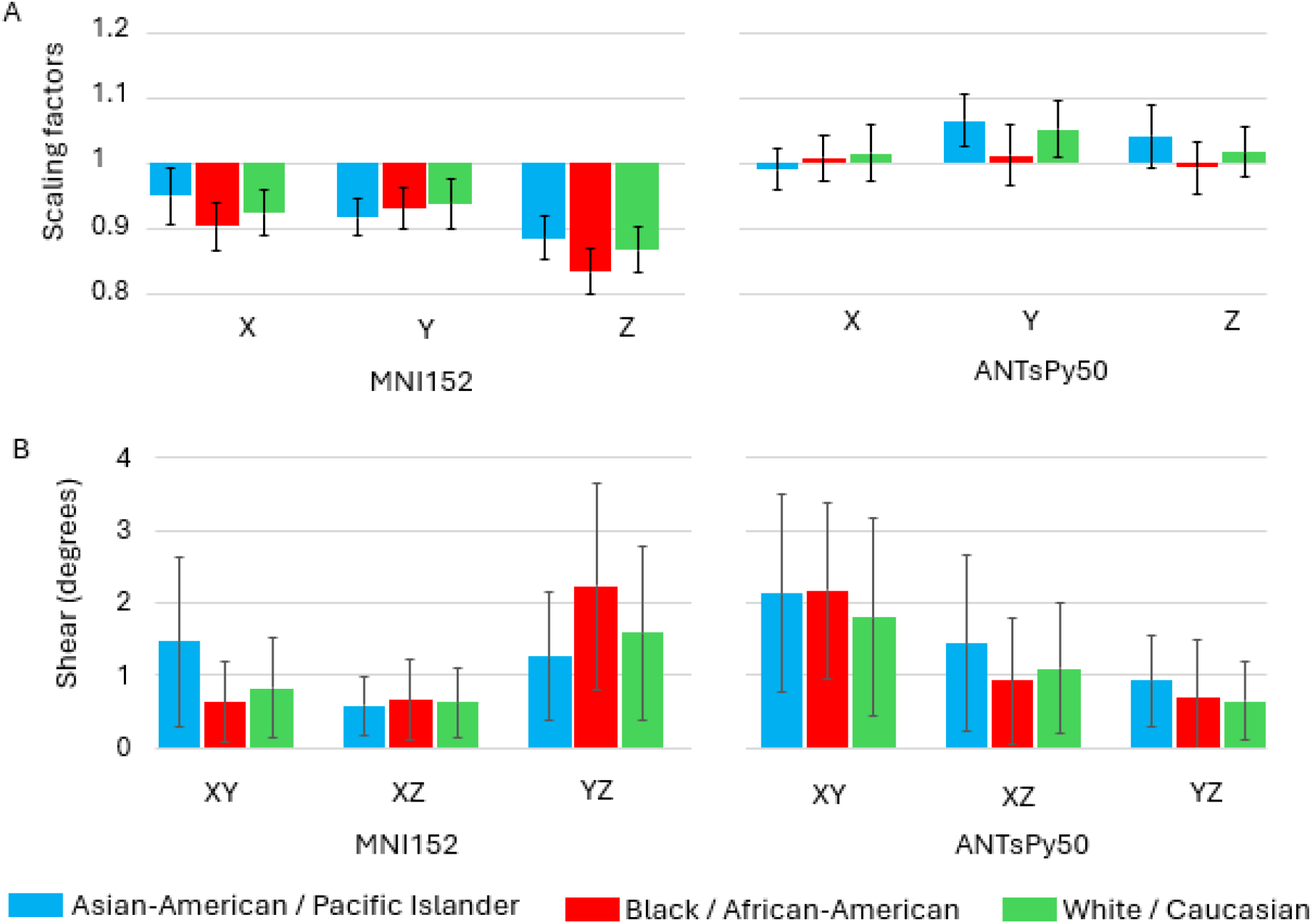
Affine transformation parameters from template-to-subject registrations for the MNI152 template and the ANTsPy50 template. (**A**) Mean scaling factors show that the MNI152 brain consistently requires contraction (scaling < 1) to match individual anatomy across all demographic groups, whereas the ANTsPy50 template requires minimal scaling, with values centered near 1.0. (**B**) Shear components, expressed in angular degrees, are small for both templates (< 2° on average), indicating that non-orthogonal distortions contribute minimally to alignment. Error bars represent the standard deviation across subjects within each demographic group.

**Figure 3.**
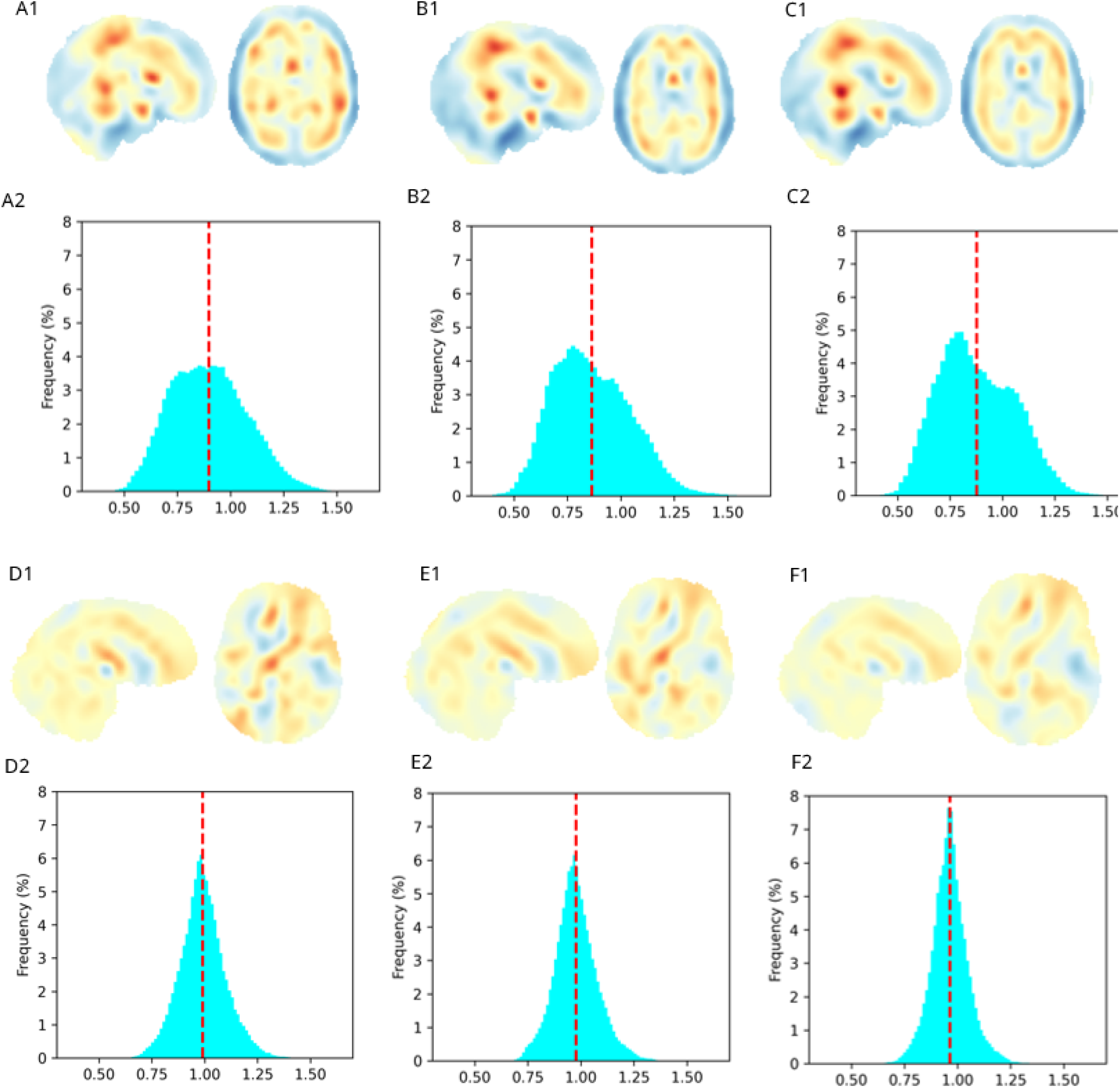
Mean Jacobian determinant (|J|) fields illustrating voxel-wise morphological differences. Representative mid-sagittal and mid-axial slices show the spatial distribution of deformations for the MNI152 brain (**A1–C1**) and the ANTsPy50 brain (**D1–F1**). Blue regions (< 1) indicate areas where the template undergoes local contraction to match subject anatomy, while red regions (> 1) indicate local expansion. Histograms for the MNI152 registration (**A2–C2**) exhibit peaks and means (dashed red line) below one, confirming a global volumetric bias, with long tails highlighting that the MNI152 discrepancies are not uniform across the brain. The ANTsPy50 registration histograms (**D2–F2**) demonstrate narrow mean deformations centered near one, indicating higher homogeneity and anatomical fidelity to the study population

### DBM

DBM analysis revealed spatially heterogeneous discrepancies between the MNI152 template and individual anatomy. The resulting |*J*| maps demonstrated that MNI152 requires systematic, nonuniform warping to align with the study population. Specifically, the temporal cortices consistently required contraction, whereas regions within the deep white matter required expansion. These patterns of deformation were highly consistent across all racial groups. Histograms of voxel-wise |*J*| values further illustrate these discrepancies, showing broad distributions with long tails extending below 0.5, (i.e., 50% local contraction). The mean |*J*| values for the MNI152 template across the entire brain were substantially below 1 (ranging from 0.87 to 0.90 for the three races), reinforcing the conclusion that the MNI152 template is systematically larger than the brains of individuals within this cohort. In contrast, the ANTsPy50 template yielded |*J*| values narrowly distributed around a mean close to 1 (ranging from 0.96 to 0.99), indicating significantly higher anatomical fidelity and minimal global bias.

## Discussion

This study provides evidence that the widely used MNI152 template does not accurately reflect the average morphology of the studied population. These findings extend prior work indicating that the MNI152 template is larger than the Talairach atlas and several race-specific templates. Crucially, our data demonstrates that this discrepancy persists even when compared directly to individual subject scans from Caucasian cohorts, suggesting the bias is an inherent artifact of the template’s construction rather than a demographic mismatch.

A key contribution of this work is the comparison between the legacy MNI152 template and an alternative template generated via ANTsPy. While the MNI152 affine scaling factors were consistently below 1.0, indicating a consistent need for contraction to match individual anatomy, the ANTsPy50 template resulted in scaling factors and mean deformations much closer to 1.0. This contrast suggests that the morphology of the MNI152 is heavily influenced by the limitations of early registration methods, such as the virtual convolution effect observed in its predecessor, the MNI305. Modern diffeomorphic averaging, as implemented in the ANTsPy pipeline, successfully produces a more representative average brain that minimizes the systematic volumetric bias. The nonlinear deformation analysis further revealed that MNI152’s discrepancies are spatially heterogeneous. Widespread contraction was required in the temporal cortices, whereas deep white matter regions often required expansion. The fact that ANTsPy50 significantly reduced these local warped errors suggests that the MNI152’s shape, not just its size, is unrepresentative of the population samples.

The systematic volumetric bias observed in MNI152 may contribute to documented inaccuracies in clinical applications, such as poor electrode-position optimization (Brahma et al., 2025). More broadly, our results highlight the importance of utilizing unbiased, population-representative template-generation pipelines. For applications requiring high anatomical fidelity, such as EEG source localization or targeted neuromodulation, the use of modern, unbiased templates or race-specific averages may provide a more accurate and reliable foundation than the MNI152 template.

